# Rapid evolutionary responses to climate change in maritime pine (*Pinus pinaster* Ait.) despite widespread adaptive lag

**DOI:** 10.64898/2026.10.14.759087

**Authors:** Thomas Francisco, Isabelle Lesur-Kupin, Ivan Scotti, Camilla Avanzi, Francesca Bagnoli, Giovanni G. Vendramin, Delphine Grivet, Marjana Westergren, Elia Vajana, Juliette Archambeau, Andrea Piotti, Santiago C. González-Martínez

## Abstract

Climate change is profoundly affecting species worldwide and poses a major threat to their long-term persistence. Although trees may have the potential for rapid adaptation to changing climates, empirical evidence remains limited, and whether these long-lived organisms can adapt quickly enough to keep pace with ongoing climate change is still an open question. To investigate real-time responses to contemporary climate change, we combined genomic data from 82 range-wide populations of the widespread tree Pinus pinaster Ait. with intensive sampling of two successive cohorts from two natural populations. Using an original genotype-environment association (GEA) approach to quantify climate (mal)adaptation, we found that most individuals from both target populations showed evidence of an adaptive lag to current climatic conditions, and are predicted to face substantial risk of maladaptation under future climates. Importantly, changes in genomic composition at climate-associated loci between cohorts were consistent with rapid evolutionary responses to ongoing climate change and could not be explained by neutral demographic processes alone, as supported by forward-time genetic simulations. We also observed a lower realised genetic load in the younger cohort, suggesting that the dynamics of deleterious mutations may change with climatic pressure. Our work provides evidence suggesting genetic responses to contemporary climate change within a single generation, including signals of rapid evolutionary changes in genomic composition at climate-associated loci and signatures of purifying selection, despite widespread adaptive lag to current climates. These findings challenge the view that trees only respond slowly to environmental change and highlight the need to integrate evolutionary dynamics into predictive models of species persistence under future climates.

## Introduction

Climate change is profoundly affecting species worldwide, posing a major threat to their long-term persistence (Parmesan 2006; Habibullah *et al*. 2022). Forest trees, owing to their long generation times and sessile nature, have been flagged as particularly at risk (Aitken *et al*. 2008), and extensive evidence of forest decline and maladaptation has already been documented (e.g., Allen *et al*. 2010; Taccoen *et al*. 2019; Browne *et al*. 2019; Benomar *et al*. 2022). *In situ* genetic adaptation, defined as changes in population allele frequencies in response to novel environmental conditions, represents one possible response of trees to changing environments (Aitken *et al*. 2008). However, although evidence of genetic adaptation is widespread in trees (reviewed in Savolainen *et al*. 2007; Leimu and Fischer 2008; Hereford 2009), documented cases of rapid adaptation (in one or a few generations) remain scarce (Saleh *et al*. 2022; Metheringham *et al*. 2025). This is particularly the case for responses to shifts in climatic baselines, raising questions about whether these long-lived organisms can adapt at the pace imposed by ongoing climate change (Aitken *et al*. 2008; Kuparinen *et al*. 2010; Fréjaville *et al*. 2020).

Evidence for evolutionary responses of trees to climate change has often been inferred from phenotypic differences among populations originating from contrasting climates, or from temporal changes in allele frequencies or phenotypic traits potentially driven by selection pressures (e.g., Jump and Peñuelas 2005; Franks *et al*. 2014; Alexandre *et al*. 2020; Gaüzere *et al*. 2020; Long *et al*. 2025). However, such studies often rely on time-consuming approaches (e.g., common gardens or long-term monitoring of natural populations) which are difficult to implement on a large scale and within the short time frames required to capture the extent and pace of contemporary climate change. Most of these approaches can document rapid responses but often lack the ability to identify fitness optima, making it difficult to assess the adaptive relevance of the observed changes (Franks *et al*. 2014). Next-generation sequencing and the emergence of research fields such as landscape genomics have offered alternative approaches for investigating climate adaptation and predicting the impacts of future climate on species (Fitzpatrick and Keller 2015; Rellstab *et al*. 2015; Capblancq *et al*. 2020). One particular approach, referred to as genomic offset, has been extensively applied to forecast future climate maladaptation, as it predicts the genomic composition required in individuals or populations to maintain current fitness under climate change (Fitzpatrick *et al*. 2021; Lachmuth *et al*. 2023; Lazic *et al*. 2024; Bonnier *et al*. 2025; Francisco *et al*. 2025; Buck *et al*. 2026; Morando-Milà *et al*. 2026). Although empirical applications remain scarce, this framework also has the potential to assess populations’ adaptive responses to climate change, for instance by comparing temporal changes in allele frequencies over generations with the direction and rate of change predicted to maintain adaptation under future climate scenarios (Dauphin *et al*. 2021).

While adaptive responses to climate change through shifts in allelic frequencies of beneficial mutations have received considerable attention, recent literature has emphasised the importance of additional processes, such as genetic load dynamics (Nadeau and Urban 2019; Aguirre-Liguori *et al*. 2021; Barratt *et al*. 2024). Indeed, genetic load, defined as the reduction in fitness caused by deleterious mutations, can increase extinction risk, particularly in populations with low effective population size (Bertorelle *et al*. 2022; Dussex *et al*. 2023; Jiang *et al*. 2025). Genetic load can be partitioned into realised load (approximated by the number of deleterious mutations in homozygosis form), which is expected to adversely affect contemporary individual fitness, and potential load (approximated by the number of deleterious mutations in heterozygosis form), which is expected to reduce future fitness if expressed as realised load. Both are mainly shaped by the interplay of genetic drift and purifying selection (Wang *et al*. 1999; Glémin 2003; Bertorelle *et al*. 2022). Quantifying the relative importance and dynamics of realised and potential genetic load is therefore crucial to better assess and predict the evolutionary responses of populations to climate change (Aguirre-Liguori *et al*. 2021; Dussex *et al*. 2023; Kremer *et al*. 2025). Furthermore, within-population variability in both climate adaptation and genetic load has rarely been explicitly considered, despite evidence that it may contribute to responses to environmental selection (e.g., Brousseau *et al*. 2013; Browne *et al*. 2019).

In this study, we combined within-population and between-cohort analyses of two contrasting *Pinus pinaster* Ait. populations to characterise (mal)adaptation to the local climate and assess single-generation genetic responses to climate change in this long-lived tree species. We analysed genomic data from two consecutive cohorts, comprising 1,481 genotyped adults and juveniles, to quantify realised and potential genetic load, as well as to assess current adaptation and future maladaptation to climate. Climate (mal)adaptation was estimated using an original genotype-environment association (GEA) approach, based on models fitted to genomic data from 82 natural populations spanning the species’ entire distribution. Finally, we tracked temporal changes in genomic composition at climate-associated loci and in genetic load between cohorts, and assessed these changes against expectations under neutral processes using neutral forward-time genetic simulations (i.e., without selection), as well as in light of observed genetic diversity and spatial genetic structure. Together, these analyses provided insights into the potential for rapid evolutionary responses of forest trees to climate change.

## Materials and Methods

### Study system

Maritime pine (*Pinus pinaster* Aiton) is a long-lived, monoecious, wind-pollinated, and anemochorous conifer tree of major economic and ecological importance (De Rigo *et al*. 2016). This species often forms large, scattered populations across southwestern Europe and North Africa, as reflected in its relatively complex population genetic structure (Bucci *et al*. 2007; Theraroz *et al*. 2024) and moderate genetic differentiation (Francisco *et al*. 2026). The species inhabits a variety of environmental zones, including Atlantic and Mediterranean coastal dunes (e.g., the Atlantic French and western Italian coasts), seasonally dry continental forests (e.g., central Spain) and Mediterranean mountainous regions (e.g., the Atlas Mountains in North Africa). Previous studies have provided evidence of local climate adaptation in *P. pinaster* populations, based on phenotypic differences among populations originating from contrasting climates (González-Martínez *et al*. 2002; de Miguel *et al*. 2022), and the identification of genomic regions potentially involved in climate adaptation (Archambeau *et al*. 2026; Francisco *et al*. 2026). Drought-induced population decline has also been documented in recent years (e.g., in some Spanish populations, Sánchez-Salguero *et al*. 2012; Gea-Izquierdo *et al*. 2019), and several areas have been flagged to be at high risk of climatic maladaptation in the future (e.g., Atlantic French populations, Theraroz *et al*. 2024; Archambeau *et al*. 2025), highlighting the potential vulnerability of *P. pinaster* to climate change.

### The two target populations

The study focused on two intensively sampled stands of *P. pinaster* located in natural forests: Lacanau State Forest (Lacanau; 44°57′48.6″ N, 1°09′53.9″ W) along the French Atlantic coast, and Tocchi State Biogenetic Natural Reserve (Tocchi; 43°07′34.2″ N, 11°15′14.2″ E) in Tuscany, Italy. These two stands (hereafter “*populations*” or “*target populations*”) differ in several aspects, primarily with regard to environmental conditions and landscape connectivity (see additional information in Theraroz 2024; Belmokhtar *et al*. 2026). Briefly, the Lacanau population is located in an Atlantic environment, characterised by relatively humid conditions, moderate temperatures, and more regular precipitation throughout the year, and is integrated within a dense network of other *P. pinaster* populations. In contrast, the Tocchi population occurs in a Mediterranean environment, characterised by hotter, drier summers and more pronounced seasonal drought, where surrounding *P. pinaster* forests are more scattered (Fig. 1).

**Figure 1.**
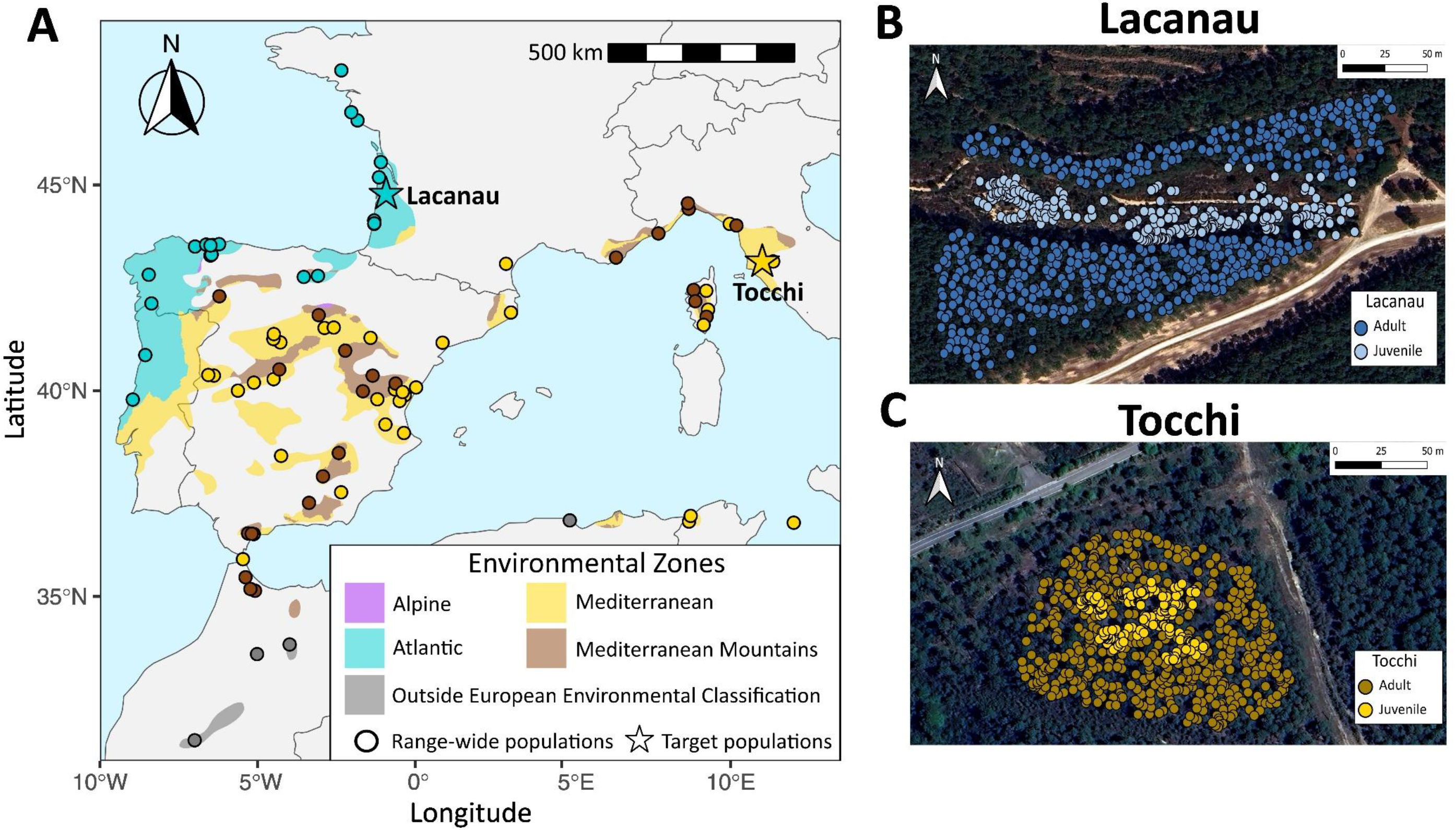
Geographical organisation of the studied populations. **A** Distribution of the 82 range-wide populations included in the study (dots), together with the two intensively sampled populations, Lacanau and Tocchi (stars), shown within the species’ range (EUFORGEN, Caudullo *et al*. 2017). Populations and the species’ range are coloured according to environmental zones defined by the European environmental zone map of Metzger (2018). **B**, **C** Orthophotos of Lacanau and Tocchi, where each dot corresponds to a genotyped tree and colours represent cohort assignment. Figure inspired from Theraroz (2024).

### Genomic datasets

Two genomic datasets were used in this study, both generated using the multispecies 4TREE Axiom array (Theraroz *et al*. 2024; Guilbaud *et al*. 2026). The first dataset, dataset 1, contained 1,518 genotyped trees from 82 natural populations spanning the species’ range of *P. pinaster*. Dataset 1 was retrieved from Francisco *et al*. (2026) after excluding the two target populations (Lacanau and Tocchi) to avoid potential over-fitting of the model used in downstream analyses. Data were filtered following Francisco *et al*. (2026) (see Supplementary Material). Further information on DNA extraction and genotyping is provided in Olsson *et al*. (2025) and Olsson *et al*. (in prep). Only SNPs shared with the second dataset, dataset 2 (see below) were retained, resulting in 8,469 SNPs. Missing data were imputed using the most common genotype within gene pools (Theraroz *et al*. 2024). Dataset 1 was used exclusively to identify climate-associated SNPs and fit the genotype-environment association (GEA) model used for genomic prediction (see below and Supplementary file 1).

Dataset 2, retrieved from Belmokhtar *et al*. (2026), consisted of 1,481 trees genotyped from the two target populations (Lacanau and Tocchi), and was used for all subsequent analyses (see Supplementary file 2). In each population, needles from 500 dominant or codominant georeferenced trees (hereafter “*adults*”) and 250 naturally regenerated georeferenced trees aged from four to sixteen years (hereafter “*juveniles*”), representing a quasi-exhaustive sampling of the investigated stands shown in Figure 1, were collected in 2021-2022. Needles were fast-dried in silica gel and DNA was extracted using the DNeasy 96 Plant Kit (Qiagen, Germany). Further information on DNA extraction and genotyping is provided in Belmokhtar *et al*. (2026). Dataset 2 was further filtered by removing both SNPs and individuals with missing data > 30% and keeping only SNPs genotyped in the 82 populations spanning the species’ range (see Fig. 1A and above), resulting in a dataset containing 8,469 SNPs genotyped in 749 trees from Lacanau and 732 trees from Tocchi. We estimated realised and potential genetic load using all the SNPs from dataset 2 located in coding regions that could be polarised (see below) and mapped (2,425 SNPs) onto a masked working version of the *Pinus tabuliformis* reference genome (Niu *et al*. 2022; Kesälahti *et al*. 2025; see Supplementary Material). For spatial genetic structure and climate (mal)adaptation analyses, missing data were imputed by the most common genotype within the target population considered. Principal component analyses (PCA) were conducted jointly on the two genomic datasets, successfully ensuring the consistency of the genomic information across datasets.

### Genetic diversity and spatial genetic structure

Genetic diversity was assessed at the individual level for Lacanau and Tocchi populations by computing individual observed heterozygosity (*H_i_*), defined as the proportion of heterozygous loci to the total number of loci per individual. Differences in *H_i_* between populations and among cohorts within populations were assessed using nested Welch’s ANOVA tests, followed by Games-Howell post-hoc pairwise comparisons.

Additionally, we characterised the fine-scale spatial genetic structure (FSGS) of these two populations by examining the spatial distribution of genetic similarity among individuals and, subsequently, estimated Wright’s neighbourhood size (*Nb*). FSGS analyses were restricted to adult trees and performed independently across populations using SPAGeDi software v1.5 (Hardy and Vekemans 2002), following De-Lucas *et al*. (2009). We quantified the decline in pairwise kinship coefficients (*F_ij_*) with increasing geographic distance across six distance classes (20, 50, 80, 100, 130, and 160 metres). The strength of FSGS (*Sp*) was then estimated as -*b*_F_/(1-*F*_1_), where *b*_F_ is the slope of the regression of kinship coefficients (*F_ij_*) on the logarithm of spatial distance and *F*_1_ corresponds to the mean kinship coefficient in the first distance class (Vekemans and Hardy 2004). *Nb* was subsequently approximated as the reciprocal of FSGS strength: *Nb*= 1/*Sp*, with higher *Nb* values suggesting a larger effective number of individuals contributing to local gene flow (Vekemans and Hardy 2004; De-Lucas *et al*. 2009; Kamdem *et al*. 2025). Analyses were replicated ten times using 500 randomly sampled SNPs and 1,000 permutations per replicate. Final *Nb* estimates were defined as the mean value across these ten replicates.

### Climatic predictors

High-resolution climatic data (30 arc-seconds) were extracted for the historical (Mid-Holocene, approximately 6,000 years ago), present (1991-2020) and future (2041-2070) periods to estimate the climate adaptive optima of the two target populations at different times (see below). Following (Francisco *et al*. 2026), additional climatic data for the 1901-1950 period (hereafter “*reference period*”), approximating the pre-Anthropocene climate conditions, were extracted to compute genotype-environment association models. The six retained climatic predictors are mean annual temperature (Annual_Tc), isothermality, temperature seasonality (Tc_seasonality), mean annual precipitation (Annual_P), precipitation seasonality (P_seasonality) and summer heat-moisture index (Summer_aridity). These predictors were selected based on their relative importance in explaining the genetic variance between populations (see Supplementary Material), as well as in relation to the ecology of the species and their importance in previous studies on the biology and local climate adaptation of *P. pinaster* (Alia and Moro 1996; Bogino and Bravo 2008; Mazza *et al*. 2014; Kurz-Besson *et al*. 2016; Caminero *et al*. 2018; Solé-Medina *et al*. 2025; Archambeau *et al*. 2026; Francisco *et al*. 2026).

Historical climate for the Mid-Holocene period were extracted from the WorldClim 1.4 database (Hijmans *et al*. 2005), while climate for the reference (1901-1950), present (1991-2020) and future (2041-2070) periods were obtained using the ClimateDT climate downscaling tool (Marchi *et al*. 2024). Climatic conditions for the historical and future periods were projected using five distinct global climate models (GCMs; Table S1). Future projections were generated under the Shared Socio-economic Pathway SSP3-7.0 (AR6), corresponding to a severe climate forcing scenario (Fujimori *et al*. 2017; Shiogama *et al*. 2023).

### Within-population variability in climate (mal)adaptation and potential for rapid evolutionary responses

#### Empirical assessment

Genotype-environment association (GEA) methods have been extensively applied to gain insights into climate (mal)adaptation across populations. These approaches identify relationships between genomic and climatic data to detect climate-associated genetic markers and can be used to project such relationships across space and time (Rellstab *et al*. 2015; Capblancq *et al*. 2020). Here, to investigate individual-level variability in climate (mal)adaptation, we propose an original two-step approach: (i) identifying climate-associated SNPs and forecasting of the optimal genomic composition under the climatic conditions experienced at the locations of the two target populations across different time periods, using a GEA model; and (ii) comparing these predicted optimal genomic compositions with the observed genomic composition at climate-associated loci across individuals from the two target populations. The first step was conducted using the range-wide dataset (dataset 1; see Fig. 1 and below). Climate-associated SNPs were identified using six complementary GEA methods (i.e., redundancy analysis and partial redundancy analysis, Capblancq and Forester 2021; latent factor mixed model, Gain and François 2021; BayPass, Gautier 2015; and gradient forest corrected or uncorrected for population genetic structure, Fitzpatrick and Keller 2015; see details in Supplementary Material). Based on previous work (Archambeau *et al*. 2026; Francisco *et al*. 2026), all SNPs identified as climate-associated candidates by any of these six methods (see details in Supplementary Material) were retained, and an RDA model was subsequently fitted using these climate-associated SNPs. This model was used for genomic forecasting to predict the optimal genomic composition under the climate of the two target populations across historical, present and future time periods (see above). To account for uncertainty in climate projections, historical and future optimum values were computed as the average across five GCMs (Table S1). In the second step, differences in observed genomic composition at climate-associated loci between cohorts within the two target populations (dataset 2) were assessed using Welch’s two-sample *t*-tests and compared to the predicted optimal genomic composition to gain insights into the potential for rapid adaptation to contemporary climate change. Under a scenario of rapid evolutionary responses, juveniles are expected to exhibit genomic compositions closer to the contemporary predicted optimum than adults.

#### Simulation evaluation

To assess whether genomic composition differences observed at climate-associated SNPs across cohorts in the two target populations reflected rapid climatic adaptation or random processes, we simulated the evolutionary dynamics of these populations under neutrality using SLiM v5.0 software (Haller *et al*. 2026). We then tested whether the observed changes deviated from neutral expectations, which would suggest the influence of adaptive processes. During the initialisation phase, 10,000 spatially georeferenced individuals were generated in a finite space, with a sex ratio of 0.5, non-overlapping generations, and pollen and seed dispersal parameters obtained from Theraroz (2024) (see Table S2 for full description of parameters). After 1,000 generations, the populations were assumed to have reached equilibrium and 500 geographically close individuals (corresponding to the “adults” in the observed data) were sampled. Then, at generation 1,001, 250 individuals (corresponding to the “juveniles” in the observed data) were sampled in the geographic area from which the 500 individuals originated.

Finally, the genotypes of the simulated individuals were retained after filtering out SNPs with a minor allele count below twelve, following the threshold applied to the empirical data. A set of SNPs was then randomly sampled to match the number of climate-associated SNPs identified (1,057 SNPs).

Mean genomic composition was subsequently compared across cohorts by projecting them onto the RDA model, using the same procedure as for the empirical data. To ensure robustness of the results, the entire simulation, including the analytical steps, was repeated 100 times for each target population (corresponding to 200 RDA axes per population).

### Realised and potential genetic load

Realised and potential genetic load were estimated to gain insight into the dynamics of deleterious mutations accumulation across Lacanau and Tocchi populations and between cohorts.

Deleteriousness of the mutations was estimated from the SNPs mapped and polarised onto the masked working version of the *Pinus tabuliformis* reference genome (Niu *et al*. 2022; see details in Supplementary Material). First SNPs causing amino-acid changes in protein-coding sequences were identified using SnpEff v5.1 software (Cingolani *et al*. 2012). Second, PROVEAN v1.1.5 software (Choi and Chan 2015) was used to predict the impact of each of these mutations on the biological function of the corresponding protein and assign a “deleteriousness” score. Following recommendations, mutations with a score lower than -2.5 were considered deleterious (Choi and Chan 2015). Finally, two genetic load metrics were computed across individuals: the realised and potential genetic load, which correspond to the number of deleterious mutations in homozygosis and heterozygosis, respectively. Differences in genetic load between populations and between cohorts within populations were assessed using Welch’s two-sample *t*-tests and nested Welch’s ANOVA, followed by Games-Howell post-hoc pairwise comparisons. As most deleterious alleles are thought to be recessive, realised load is expected to have a stronger immediate impact on fitness, whereas potential load can be converted into realised load in future generations (González-Martínez *et al*. 2017; Dussex *et al*. 2023).

## Results

### Genetic diversity and spatial genetic structure

Individual observed heterozygosity (*H_i_*) differed both between populations and between cohorts within populations. Overall, *H_i_* was larger in Lacanau (0.249) than in Tocchi (0.194), a pattern further confirmed at the cohort-level (Table 1 and Fig. S1; see statistical tests in Table S3). Within each population, juveniles, on average, consistently displayed higher *H_i_* than adults.

**Table 1.** Population characteristics. Descriptive statistics for Lacanau and Tocchi populations, including the number of successfully genotyped individuals (N), individual observed heterozygosity (*H_i_*) and Wright’s neighbourhood size (*Nb*; see full description in Table S4). Post hoc pairwise comparisons from one-way Welch’s ANOVA using Games-Howell tests are shown for *H_i_* with lowercase letters (*p*-value < 0.05; see Table S3 for details).

| Populations | Cohorts | *N* | *H<sub>i</sub>* | *Nb* |
| --- | --- | --- | --- | --- |
| Lacanau | Adult | 500 | 0.24a | 662 |
|  | Juvenile | 249 | 0.27b |  |
| Tocchi | Adult | 484 | 0.19c | 106 |
|  | Juvenile | 248 | 0.20d |  |

Spatial genetic structure also diverged markedly between populations, with neighbourhood size (*Nb*) being over six times higher in Lacanau than in Tocchi (Table 1 and S4). Altogether, these results suggest larger local effective population size and connectivity in Lacanau than in Tocchi, as well as increased genetic diversity in juvenile cohorts in both populations.

### Within-population variability in climate (mal)adaptation variability within populations and potential for rapid evolutionary responses

#### Climate-associated SNPs and RDA model

Re-analysis of the dataset 1 from Francisco *et al*. (2026), comprising 82 populations after excluding Lacanau and Tocchi populations, and the six retained climatic predictors identified 1,057 climate-associated SNPs across the six GEA analyses (Fig. S2). The RDA model subsequently fitted using these SNPs explained 71.1% of the genetic variance among the 82 populations. The first axis (RDA1) captured variation in precipitation and temperature seasonality, as well as summer aridity (negative loadings on RDA1) and isothermality (positive loadings on RDA1; Fig. 2). The second axis (RDA2) was mainly driven by summer aridity, isothermality and precipitation seasonality (positive loadings on RDA2), and more marginally by annual temperature and precipitation (negative loadings on RDA2).

**Figure 2.**
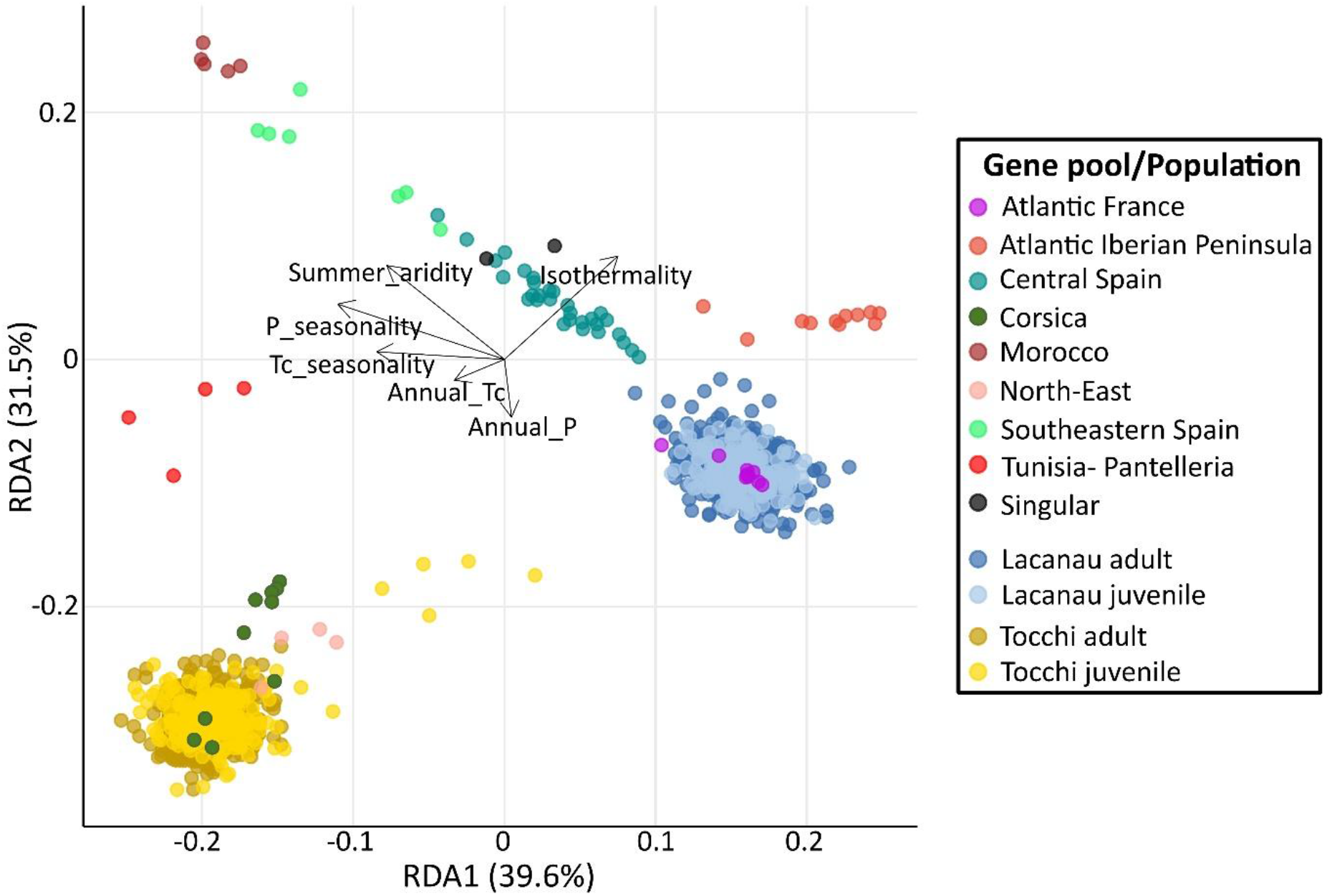
Description of the RDA model used to infer climate (mal)adaptation. Biplot of the RDA model showing the relative contribution of the climatic predictors to the first two RDA axes and the distribution of the populations used to fit the model (see Fig. S3 for a genotype-level representation). Genotyped individuals from Lacanau and Tocchi populations are projected into this space based on their genomic composition at climate-associated loci. Dots are coloured according to gene pool (Theraroz *et al*. 2024) or population cohort assignment. To facilitate readability, distinct gene pools consisting of a single population have been grouped under the term “*singular*” (see details in Theraroz *et al*. 2024).

Projecting individuals from Lacanau and Tocchi onto the RDA space first revealed major differences in climate-associated SNP frequencies (RDA values) between the two populations (Fig. 2). Lacanau trees clustered in the middle-to-lower right part of the RDA space, sharing RDA values with other Atlantic French populations, and suggesting association with high annual precipitation, as well as low summer aridity, precipitation and temperature seasonality. In contrast, Tocchi trees were located in the bottom-left part of the RDA space, close to other geographically distant Mediterranean populations from Corsica and Italy, and appeared associated with high annual temperature, precipitation and temperature seasonality, as well as lower isothermality. The correspondence between population positions, the climatic conditions of their native environments, and the position of populations from similar environments suggests that the RDA model successfully captured climate-driven allelic frequency patterns.

#### Variability in climate (mal)adaptation within populations and across cohorts

We found substantial within-population variability in genomic composition at climate-associated loci (RDA values) in both the Lacanau and Tocchi populations, suggesting individual-level variation in climate adaptation (Fig. 2). The magnitude of this variation was comparable to that observed across range-wide populations in dataset 1 (Fig. S3). Density plots confirmed this high variability, with a broad-range of RDA values and, in Tocchi, a long distribution tail along both RDA axes (Fig. 3). We further found that almost all trees in both populations were markedly distant from the predicted genomic optima along both RDA axes, with the exception of five juveniles in Tocchi (i.e., ITA00019-540, ITA00019-551, ITA00019-563, ITA00019-611 and ITA00019-734). However, Lacanau and Tocchi trees differed in their positions relative to these predicted optima. Lacanau trees were located between the historical and present/future optima, with trees being closer to the present/future optima along RDA1 and to the historical optimum along RDA2. Conversely, most Tocchi trees were located beyond the present/future optima, with genomic composition at climate-associated loci closer to the present/future optima than to the historical optimum along both RDA axes.

**Figure 3.**
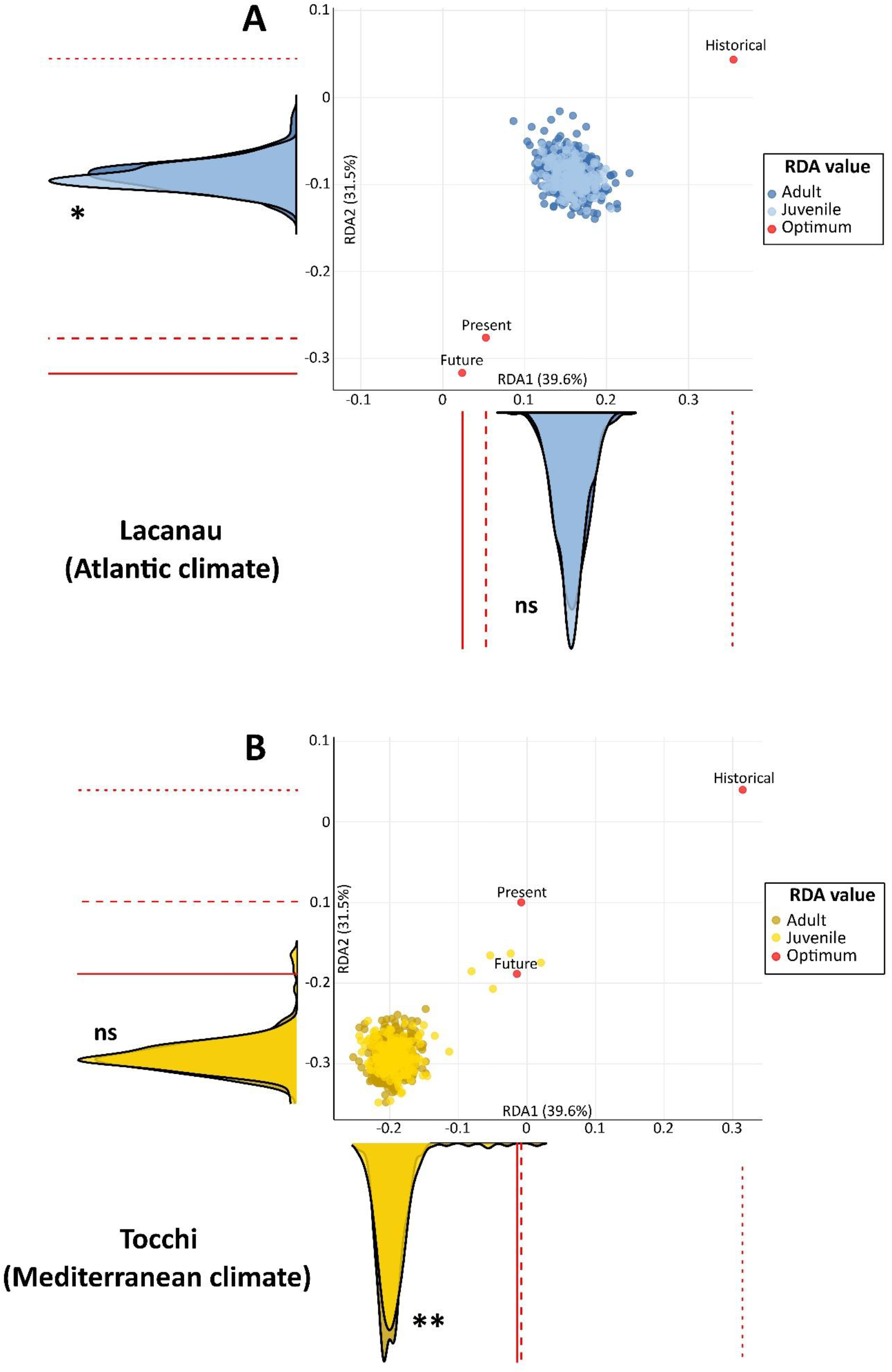
Within-population variability in climate (mal)adaptation and potential for rapid evolutionary responses. Focus on the projected positions of genotyped individuals from (**A**) Lacanau and (**B**) Tocchi within the RDA model presented in Fig. 2. Individuals are positioned according to their genomic composition at climate-associated loci and coloured by cohort, expressed as RDA scores. Predicted genomic composition optima, also expressed as RDA scores, are shown for historical (Mid-Holocene, approximately 6,000 years ago), present (1991-2020) and future (2041-2070) time periods. Density plots show cohort-specific distributions of RDA scores, with differences between cohorts assessed using Welch’s two-sample *t*-tests (^ns^ not significant; *: *p*-value < 0.05; **: *p*-value < 0.01). Optimum values are also represented as lines on the density plots, corresponding to historical (dotted), present (dashed), and future (solid) time periods.

Interestingly, despite differences in population positions relative to the optima, comparison of genomic composition at climate-associated loci between cohorts within populations, as shown by RDA, revealed similar patterns in Lacanau and Tocchi (Fig. 3). In Lacanau, no significant shifts in RDA values were observed between adults and juveniles along RDA1, while juveniles were significantly closer to the present and future optima than adults along RDA2. Similarly, in Tocchi, juveniles were significantly closer to the present and future optimum values than adults along RDA1, although no significant difference between adults and juveniles were found along RDA2 (see details in Table S3).

#### Simulation evaluation

Neutral simulations (i.e., without selection) showed that shifts in genomic composition at climate-associated loci between adults and juveniles comparable, to those observed empirically, occurred in only six out of 200 simulated axes (3%) for Lacanau and in eight out of 200 (4%) for Tocchi, regardless of shift direction. When considering only shifts in the same direction as observed empirically (i.e., with juveniles closer to the predicted genomic optima than adults), such shifts occurred in only 1.5% of simulated axes, supporting that the observed cohort differences were rarely reproduced under the neutral simulations.

### Realised and potential genetic load

Comparison of realised and potential load estimates revealed contrasting patterns between populations, with Lacanau trees exhibiting, on average, a significantly lower number of deleterious mutations in homozygosis, but a higher number in heterozygosis compared to Tocchi (median of 90 vs. 107 deleterious mutations in homozygosis and 97 vs. 66 in heterozygosis, respectively; Fig. 4). In both populations, realised and potential load were negatively correlated (Pearson’s correlations of - 0.69 for Lacanau and -0.80 for Tocchi).

**Figure 4.**
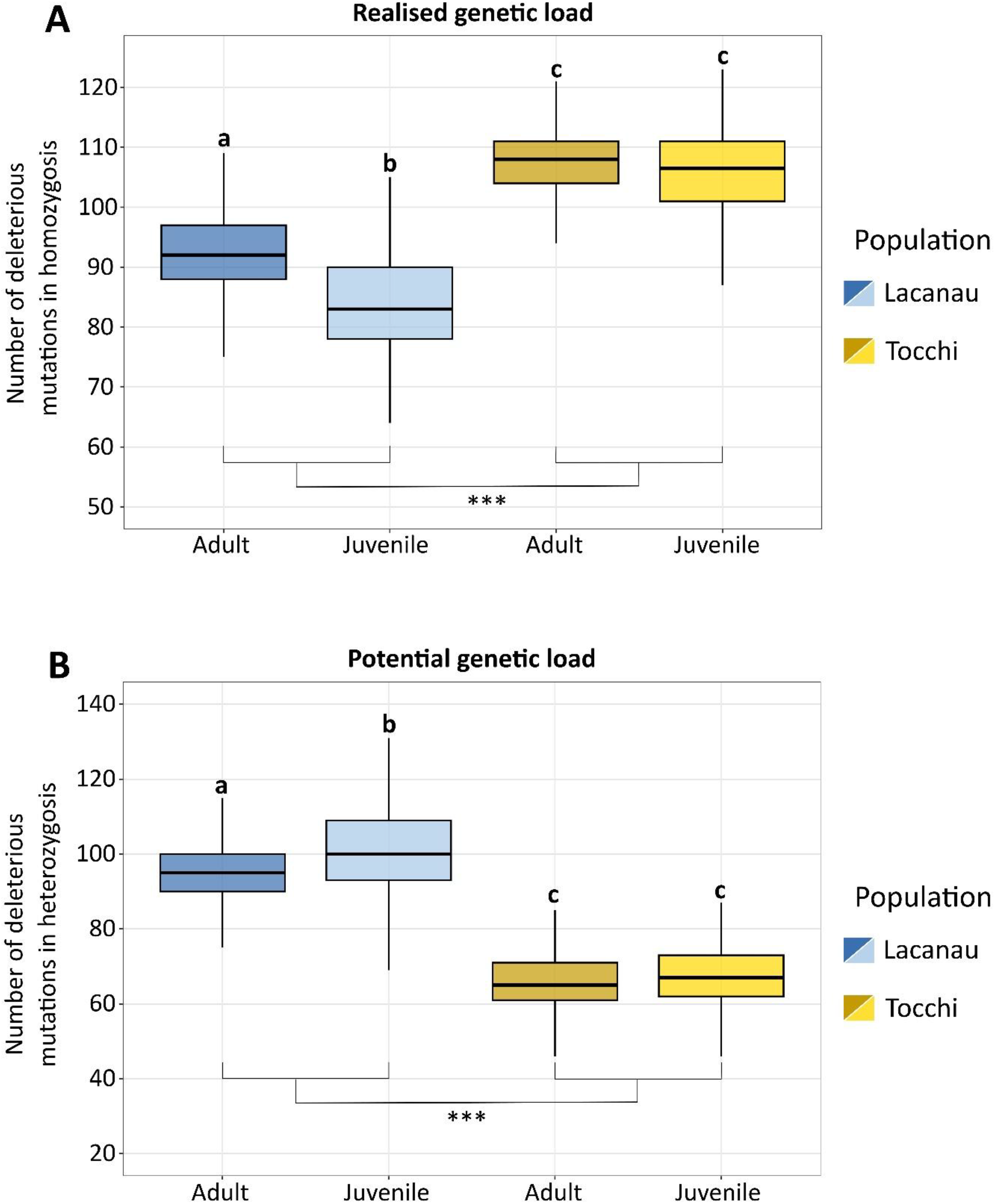
Genetic load. Comparisons of realised (**A**) and potential (**B**) genetic load across populations and cohorts. Games-Howell’s post hoc groups are shown by lowercase letters (*p*-value < 0.05). Pairwise population differences were tested using Welch’s two-sample *t*-tests (***: *p*-value < 0.001).

Additionally, differences in dynamics were also observed between cohorts across populations (see Fig. 4 and Table S3). In Lacanau, significant changes in the number of deleterious mutations were observed between cohorts, with juveniles exhibiting a lower number of deleterious mutations in homozygosis than adults (median values in juveniles equal to 83 vs. 92 in adults) but a higher number in heterozygosis (median values of 100 vs. 95). In contrast, no significant changes between cohorts were observed in the Tocchi population. This pattern could suggest ongoing purging of realised genetic load in Lacanau whereas no comparable evidence of purging was detected in Tocchi.

## Discussion

### Extensive within-population variation at climate-associated loci and evidence for adaptation lags

Substantial individual-level variation in climate adaptation was suggested within both the Lacanau and Tocchi populations, irrespective of the cohorts considered, with the dispersion in RDA space being of a similar magnitude to that observed among population means within gene pools across the range-wide populations (dataset 1). However, comparisons between target and range-wide populations should be interpreted with caution due to the unbalanced number of genotyped individuals across populations (i.e., much higher in the two target populations). These results support the idea that both target populations harbour large standing adaptive genetic variation, which may promote rapid genetic adaptation to changing environments (Barrett and Schluter 2008; Messer and Petrov 2013; Matuszewski *et al*. 2015). However, nearly all individuals were genetically distant from the genomic composition predicted to be optimal under the present-day climate, suggesting a pronounced adaptation lag to contemporary climate. Climatic changes since the Mid-Holocene (i.e., historical period; ∼6,000 years ago) could explain the apparent adaptive lag observed in Lacanau.

Indeed, Lacanau individuals exhibited genomic compositions at climate-associated loci that were intermediate between those predicted for the Mid-Holocene and the present day, a pattern further supported by their position relative to the reference time period (1901-1950; Fig. S4). This suggests that Lacanau individuals are experiencing an adaptive lag in response to climatic changes that have occurred since the Mid-Holocene. Comparable adaptive lags have been documented in several tree species (Browne *et al*. 2019; Fréjaville *et al*. 2020; Goetz *et al*. 2026), suggesting that long-term maladaptation may be more widespread than currently assumed and highlighting the need for additional case studies across forest tree species. Remarkably, genomic composition at climate-associated loci in Tocchi did not correspond to an intermediate state between historical and current optima and were instead associated with more extreme precipitation and temperature seasonality, as well as reduced isothermality. This pattern may reflect the legacy of post-glacial recolonisation from Last Glacial Maximum (LGM) refugia. Indeed, several studies suggested that northern Italian populations, including Tocchi, originated from southern Italian or eastern North African refugial regions following the LGM, as opposed to Atlantic French populations which likely persisted locally along the French Atlantic coast (Vendramin *et al*. 1998; Bucci *et al*. 2007). Consequently, contemporary Tocchi trees may have retained adaptive genomic signatures inherited from ancestral climatic conditions experienced in these refugial areas, which could explain the currently observed patterns.

Overall, these results suggest that, despite substantial within-population variability at climate-associated loci and a likely distinct post-glacial history, both Lacanau and Tocchi populations share patterns consistent with extensive adaptive lags to contemporary climate. Furthermore, adaptive lags are projected to intensify under future climatic conditions, with most individuals predicted to lie farther from future than current genomic optima, suggesting substantial maladaptation to future climate in the absence of genetic responses.

### Contrasting dynamics of realised and potential genetic load between cohorts and among populations

Realised (homozygous) and potential (heterozygous) genetic load showed a negative relationship, mirroring theoretical expectations (Bertorelle *et al*. 2022). Their intrinsic relationship can explain the observed pattern, where potential and realised loads may convert into one another due to various evolutionary processes, including genetic drift and purifying selection (Mathur and DeWoody 2021; Dussex *et al*. 2023). Genetic load dynamics varied across populations, with Lacanau exhibiting lower realised load, and consequently higher potential load, than Tocchi. The Lacanau population also displayed a significant reduction of realised load and an increase of potential load in juveniles compared to adults, suggesting ongoing purifying selection, while no such differences between cohorts were observed in Tocchi. Differences in the intensity of genetic drift may explain these patterns, as Tocchi showed both lower genetic diversity (*H_i_*) and a smaller effective number of individuals contributing to local gene flow (*Nb*) than Lacanau, potentially reflecting and enhancing stronger genetic drift, respectively. Stronger genetic drift is expected to reduce the efficiency of purifying selection, which can result in higher fixation of deleterious alleles (i.e., potential load converted into realised load), as previously observed in various studies (Jaramillo-Correa *et al*. 2020; Hasselgren *et al*. 2021; Dussex *et al*. 2021, 2023). Overall, these results suggest contrasting levels and dynamics of genetic load and provide evidence that selection during the juvenile stage may reduce realised genetic load in trees under certain conditions, as suggested by Kremer *et al*. (2025).

However, other processes, including sampling effects or stochastic mortality, could also have contributed to the observed differences in genetic load between cohorts and populations. The persistence of these populations under climate change may be impacted differentially by the different kinds of genetic load. Tocchi is likely to exhibit a reduced capacity for adaptive genetic responses, whereas Lacanau may face a greater risk of future fitness decline owing to its higher potential genetic load, particularly under adverse demographic scenarios such as population decline or bottlenecks.

### Rapid evolutionary responses to climate change at climate-associated loci

Rapid evolutionary responses to ongoing climate change are suggested by the observed shifts in genomic composition at climate-associated loci between successive cohorts, with juveniles being significantly closer than adults to the present-day and, subsequently, future, predicted optimal genomic compositions. Our simulation approach further suggests that these patterns are unlikely to be explained by neutral demographic processes alone. Shifts across cohorts were observed in both Lacanau and Tocchi populations, but involved different climatic predictors and occurred only along the RDA axes for which the genomic compositions at climate-associated loci were the furthest from the present optimum (i.e., in cases with higher adaptation lag and presumably stronger on-going selection). Therefore, historical adaptation lag in these populations appears to enhance current genetic adaptation by potentially imposing stronger selection pressure on the trees. Consistent with these findings, Theraroz (2024) reported evidence of contemporary directional selection acting on growth and functional traits in the same two *P. pinaster* populations. Overall, our work contributes to the still limited body of evidence documenting rapid genetic responses of forest trees to climatic changes (Saleh *et al*. 2022; Metheringham *et al*. 2025; Eberhardt *et al*. 2026), using an original genomic approach based on genotype-environment association.

Interestingly, a few unrelated juveniles in Tocchi, as revealed by kinship analysis, were located very close to both the present-day and future predicted genomic composition optima, suggesting that they may be nearly locally adapted to contemporary climate and even pre-adapted to future conditions along these climatic predictors. Combined with evidence supporting the presence of substantial standing genetic variation at climate-associated loci and the significant shifts observed between cohorts, these results highlight the importance of within-population variation in shaping both current and future adaptive responses to climate change. They also underscore the need to integrate such variation more systematically into predictive models, as suggested by previous literature (e.g., Savolainen *et al*. 2007; Brousseau *et al*. 2013; Scotti *et al*. 2016; Browne *et al*. 2019). Nevertheless, it remains unclear whether these few individuals will contribute to accelerating population-level genetic adaptation to changing climates, or instead be lost through stochastic processes, or antagonistic selection imposed by other abiotic or biotic factors (Kirkpatrick and Peischl 2013; Orr and Unckless 2014; Charlesworth 2020; Aitken *et al*. 2024).

Most trees exhibited evidence of adaptive lag, suggesting that climate change may still be occurring at rates challenging the capacity for genetic adaptation, potentially leading to population declines and local extinctions (Shaw and Etterson 2012). Similar conclusions have been drawn for several populations of *Pinus cembra* in the Swiss Alps (Dauphin *et al*. 2021), raising concerns that such patterns may be widespread among organisms with long-generation like trees. Nevertheless, these conclusions should be interpreted with caution, as several sources of uncertainty may influence the results. These include uncertainties associated with historical climate reconstructions, future climate projections, and the choice of emission scenarios (e.g., Braconnot *et al*. 2012; Hausfather *et al*. 2020; Zelinka *et al*. 2020), as well as uncertainties inherent to genotype-environment association models via omission of relevant climatic predictors and genomic variants (e.g., structural variants) or failure to identify adaptive loci displaying complex adaptive pathways (e.g., conditional neutrality; Yoder and Tiffin 2018). It also remains to be determined whether the observed shifts in genomic composition at climate-associated loci between successive cohorts translate into fitness advantages for younger trees. Finally, additional adaptive mechanisms, including phenotypic plasticity, may help mitigate the negative impacts of climate change on these populations.

## Conclusion

We present an original approach using genotype-environment associations to investigate short-term genetic responses of forest tree populations to climatic change, and to quantify their current adaptive lags. Our work revealed substantial variation at climate-associated loci, suggesting the presence of extensive standing adaptive genetic variation within populations. We also found evidence suggesting (i) rapid evolutionary responses to climate change, as reflected in changes in genomic composition at climate-associated loci, and (ii) purifying selection acting on realised genetic load across cohorts, challenging the view that trees only respond slowly to environmental change. Nonetheless, widespread adaptation lags to contemporary climate, and its projected intensification under future climatic conditions, suggests that climate may still be changing at rates that exceed the capacity of tree populations for genetic adaptation. This highlights the need for further studies across forest tree species to better understand the pace and limits of climate adaptation. In particular, determining whether the dynamics of genetic load are changing in response to climate change, and assessing the extent to which phenotypic plasticity, among other adaptive processes, contribute to observed patterns of climate adaptation in trees, will be critical for developing robust forecasts of forest tree responses in a rapidly changing world.

## Author Contributions

Designed research: TF, AP, SCG-M, MW, JA; acquired and pretreated genomic data: CA, FB, EV, GGV, SO, DG; analysed data: TF, IL-K, SCG-M, IS; wrote the paper: TF, SCG-M; obtained funding: GGV, DG, SCG-M, MW, IS. All authors participated in discussions and revised multiple versions of the manuscript, including the final version.

## Acknowledgments

TF was funded by a PhD grant of the INRAE Scientific Department ECODIV. EV was supported by the MSCA European fellowship MedForAct (GA 101107604). We thank the INRAE - UEFP Forest Experimental Facility, Cestas, France (https://doi.org/10.15454/1.5483264699193726E12), for assistance with field work. This research was funded by EU H2020 FORGENIUS (grant agreement No. 862221) and Horizon Europe OptFORESTS (grant agreement No. 101081774) projects. This work was also supported by the French National Research Agency (ANR) under the France 2030 program (project MICROFOREST, ANR-25-PEFO-0007). Views and opinions expressed are however those of the authors only and do not necessarily reflect those of the European Union. Neither the European Union nor the granting authority can be held responsible for them. We are grateful to the Genotoul bioinformatics platform Toulouse Occitanie (Bioinfo Genotoul, https://bioinfo.genotoul.fr/) for providing computing and storage resources. TF thanks the EVOLTREE network (https://www.evoltree.eu/) for funding a travel grant to attend a workshop on forward-time genetic simulations.

## Data Archiving Statement

Genomic data for the 82 range-wide populations from Olsson *et al*. (2025) and Francisco *et al*. (2026) is available at Zenodo (https://doi.org/10.5281/zenodo.14950394). Genomic data for the two target populations, Lacanau and Tocchi, from Belmokhtar *et al*. (2026) is available at Zenodo (https://doi.org/10.5281/zenodo.17054530).

## Code availability

The code used for data processing and analyses is publicly available at the following GitHub repository: https://github.com/Thomas-Francisco/Rapid-evolutionary-responses-to-contemporary-climate-change-in-Pinus-pinaster-despite-adaptive-lag.

## References

Aguirre-Liguori JA, Ramírez-Barahona S, Gaut BS (2021) The evolutionary genomics of species’ responses to climate change. Nature Ecology & Evolution 5:1350–1360.

Aitken SN, Jordan R, Tumas HR (2024) Conserving Evolutionary Potential: Combining Landscape Genomics with Established Methods to Inform Plant Conservation. Annual Review of Plant Biology 75:707–736.

Aitken SN, Yeaman S, Holliday JA, et al. (2008) Adaptation, migration or extirpation: climate change outcomes for tree populations. Evolutionary Applications 1:95–111.

Alexandre H, Truffaut L, Klein E, et al. (2020) How does contemporary selection shape oak phenotypes? Evolutionary Applications 13:2772–2790.

Alia R, Moro J (1996) Performance of *Pinus pinaster* Ait. provenances in Central Spain. Forest Systems 5:57–75.

Allen CD, Macalady AK, Chenchouni H, et al. (2010) A global overview of drought and heat-induced tree mortality reveals emerging climate change risks for forests. Forest Ecology and Management 259:660–684.

Archambeau J, Benito GM, de MM, et al (2026) Evaluating Genomic Offset Predictions in a Forest Tree with High Population Genetic Structure. The American Naturalist 207:389–414.

Barratt CD, Onstein RE, Pinsky ML, et al. (2024) Life on the edge: A new toolbox for population-level climate change vulnerability assessments. Methods in Ecology and Evolution 15:2038–2058.

Barrett RDH, Schluter D (2008) Adaptation from standing genetic variation. Trends in Ecology & Evolution 23:38–44.

Belmokhtar N, Mestre Runge C, Avanzi C, et al. (2026) An integrative set of high-throughput spectroscopic, phenotypic, genetic and remote sensing environmental data: 1. Populus nigra and Pinus pinaster.

Benomar L, Bousquet J, Perron M, et al. (2022) Tree Maladaptation Under Mid-Latitude Early Spring Warming and Late Cold Spell: Implications for Assisted Migration. Frontiers in Plant Science 13.

Bertorelle G, Raffini F, Bosse M, et al. (2022) Genetic load: genomic estimates and applications in non-model animals. Nature Reviews Genetics 23:492–503.

Bogino SM, Bravo F (2008) Growth response of *Pinus pinaster* Ait. to climatic variables in central Spanish forests. Annals of Forest Science 65:506–506.

Bonnier J, Sáez Laguna E, Francisco T, et al. (2025) Wet Season Environments Drive Local Adaptation in the Timber Tree *Dicorynia guianensis* in French Guiana. Molecular Ecology e17759.

Braconnot P, Harrison SP, Kageyama M, et al. (2012) Evaluation of climate models using palaeoclimatic data. Nature Clim Change 2:417–424.

Brousseau L, Bonal D, Cigna J, Scotti I (2013) Highly local environmental variability promotes intrapopulation divergence of quantitative traits: an example from tropical rain forest trees. Annals of Botany 112:1169–1179.

Browne L, Wright JW, Fitz-Gibbon S, et al. (2019) Adaptational lag to temperature in valley oak (*Quercus lobata*) can be mitigated by genome-informed assisted gene flow. Proceedings of the National Academy of Sciences USA 116:25179–25185.

Bucci G, González-Martínez SC, Le Provost G, et al. (2007) Range-wide phylogeography and gene zones in *Pinus pinaster* Ait. revealed by chloroplast microsatellite markers. Molecular Ecology 16:2137–2153.

Buck RC, Butterfield HS, Hiroyasu EHT, et al. (2026) Applying Both Landscape Genomic and Ecological Niche Model Predictions to Inform Conservation Strategies of a California Foundational Oak Species. Molecular Ecology 35:e70322.

Caminero L, Génova M, Camarero JJ, Sánchez-Salguero R (2018) Growth responses to climate and drought at the southernmost European limit of Mediterranean *Pinus pinaster* forests. Dendrochronologia 48:20–29.

Capblancq T, Fitzpatrick MC, Bay RA, et al. (2020) Genomic Prediction of (Mal)Adaptation Across Current and Future Climatic Landscapes. Annual Review of Ecology, Evolution, and Systematics 51:245–269.

Capblancq T, Forester BR (2021) Redundancy analysis: A Swiss Army Knife for landscape genomics. Methods in Ecology and Evolution 12:2298–2309.

Caudullo G, Welk E, San-Miguel-Ayanz J (2017) Chorological maps for the main European woody species. Data in Brief 12:662–666.

Charlesworth B (2020) How Long Does It Take to Fix a Favorable Mutation, and Why Should We Care? The American Naturalist 195:753–771.

Choi Y, Chan AP (2015) PROVEAN web server: a tool to predict the functional effect of amino acid substitutions and indels. Bioinformatics 31:2745–2747.

Cingolani P, Platts A, Wang LL, et al. (2012) A program for annotating and predicting the effects of single nucleotide polymorphisms, SnpEff. Fly 6:80–92.

Dauphin B, Rellstab C, Schmid M, et al. (2021) Genomic vulnerability to rapid climate warming in a tree species with a long generation time. Global Change Biology 27:1181–1195.

de Miguel M, Rodríguez-Quilón I, Heuertz M, et al. (2022) Polygenic adaptation and negative selection across traits, years and environments in a long-lived plant species (*Pinus pinaster* Ait., Pinaceae). Molecular Ecology 31:2089–2105.

De Rigo D, Caudullo G, Durrant TH, et al. (2016) European Atlas of Forest Tree Species. LUX

Dussex N, Morales HE, Grossen C, et al. (2023) Purging and accumulation of genetic load in conservation. Trends in Ecology & Evolution 38:961–969.

Dussex N, van der Valk T, Morales HE, et al. (2021) Population genomics of the critically endangered kākāpō. Cell Genomics 1:100002.

Eberhardt L, Reuss F, Blázquez MEN, et al. (2026) Climate change intensifies rapid genomic selection beyond the ancestral niche of Fagus sylvatica. 2026.03.09.710448

Fitzpatrick MC, Chhatre VE, Soolanayakanahally RY, Keller SR (2021) Experimental support for genomic prediction of climate maladaptation using the machine learning approach Gradient Forests. Molecular Ecology Resources 21:2749–2765.

Fitzpatrick MC, Keller SR (2015) Ecological genomics meets community-level modelling of biodiversity: mapping the genomic landscape of current and future environmental adaptation. Ecology Letters 18:1–16.

Francisco T, Lesur-Kupin I, Guadaño-Peyrot C, et al. (2026) Demographic history shapes forest tree vulnerability to climate change. 2026.03.10.710859

Francisco T, Mayol M, Vajana E, et al. (2025) Genomic Signatures of Climate-Driven (Mal)Adaptation in an Iconic Conifer, the English Yew (*Taxus baccata* L.). Evolutionary Applications 18:e70160.

Franks SJ, Weber JJ, Aitken SN (2014) Evolutionary and plastic responses to climate change in terrestrial plant populations. Evolutionary Applications 7:123–139.

Fréjaville T, Vizcaíno-Palomar N, Fady B, et al. (2020) Range margin populations show high climate adaptation lags in European trees. Global Change Biology 26:484–495.

Fujimori S, Hasegawa T, Masui T, et al. (2017) SSP3: AIM implementation of Shared Socioeconomic Pathways. Global Environmental Change 42:268–283.

Gain C, François O (2021) LEA 3: Factor models in population genetics and ecological genomics with R. Molecular Ecology Resources 21:2738–2748.

Gautier M (2015) Genome-Wide Scan for Adaptive Divergence and Association with Population-Specific Covariates. Genetics 201:1555–1579.

Gaüzere J, Klein EK, Brendel O, et al. (2020) Microgeographic adaptation and the effect of pollen flow on the adaptive potential of a temperate tree species. New Phytologist 227:641–653.

Gea-Izquierdo G, Férriz M, García-Garrido S, et al. (2019) Synergistic abiotic and biotic stressors explain widespread decline of *Pinus pinaster* in a mixed forest. Science of The Total Environment 685:963–975.

Glémin S (2003) How are deleterious mutations purged? Drift versus nonrandom mating. Evolution 57:2678–2687.

Goetz ARB, Ochoa ME, Badillo B, et al. (2026) Persistent climate maladaptation decreases performance of a keystone tree species (Quercus lobata). 2026.02.06.704494

González-Martínez SC, Alía R, Gil L (2002) Population genetic structure in a Mediterranean pine (*Pinus pinaster* Ait.): a comparison of allozyme markers and quantitative traits. Heredity 89:199–206.

González-Martínez SC, Ridout K, Pannell JR (2017) Range Expansion Compromises Adaptive Evolution in an Outcrossing Plant. Current Biology 27:2544–2551.e4.

Guilbaud R, Bagnoli F, Ben-Sadoun S, et al. (2026) Development of the 4TREE SNP array, a forest multispecies array to enhance European Breeding and conservation programs in pine, poplar and ash. 2026.03.21.711309

Habibullah MS, Din BH, Tan S-H, Zahid H (2022) Impact of climate change on biodiversity loss: global evidence. Environmental Science and Pollution Research 29:1073–1086.

Haller BC, Ralph PL, Messer PW (2026) SLiM 5: Eco-evolutionary Simulations Across Multiple Chromosomes and Full Genomes. Molecular Biology and Evolution 43:msaf313.

Hardy OJ, Vekemans X (2002) spagedi: a versatile computer program to analyse spatial genetic structure at the individual or population levels. Molecular Ecology Notes 2:618–620.

Hasselgren M, Dussex N, von Seth J, et al. (2021) Genomic and fitness consequences of inbreeding in an endangered carnivore. Molecular Ecology 30:2790–2799.

Hausfather Z, Drake HF, Abbott T, Schmidt GA (2020) Evaluating the Performance of Past Climate Model Projections. Geophysical Research Letters 47:e2019GL085378.

Hereford J (2009) A quantitative survey of local adaptation and fitness trade-offs. The American Naturalist 173:579–588.

Hijmans RJ, Cameron SE, Parra JL, et al. (2005) Very high resolution interpolated climate surfaces for global land areas. International Journal of Climatology 25:1965–1978.

Jaramillo-Correa JP, Bagnoli F, Grivet D, et al. (2020) Evolutionary rate and genetic load in an emblematic Mediterranean tree following an ancient and prolonged population collapse. Molecular Ecology 29:4797–4811.

Jiang J, Chen J-F, Li X-T, et al. (2025) Incorporating genetic load contributes to predicting *Arabidopsis thaliana*’s response to climate change. Nature Communications 16:2752.

Jump AS, Peñuelas J (2005) Running to stand still: adaptation and the response of plants to rapid climate change. Ecology Letters 8:1010–1020.

Kamdem NG, Sonké B, Sergeant S, et al. (2025) Short-distance seed and pollen dispersal in both hunted and intact forests in the lower canopy African rainforest tree, Coula edulis Baill. (Coulaceae). BMC Ecology and Evolution 25:20.

Kesälahti R, Kumpula TA, Cervantes S, et al. (2025) Optimising Exome Captures in Species With Large Genomes Using Species-Specific Repetitive DNA Blocker. Molecular Ecology Resources 25:e14053.

Kirkpatrick M, Peischl S (2013) Evolutionary rescue by beneficial mutations in environments that change in space and time. Philosophical Transactions B 368:20120082.

Kremer A, Chen J, Lascoux M (2025) ‘Chimes of resilience’: what makes forest trees genetically resilient? New Phytologist 246:1934–1951.

Kuparinen A, Savolainen O, Schurr FM (2010) Increased mortality can promote evolutionary adaptation of forest trees to climate change. Forest Ecology and Management 259:1003–1008.

Kurz-Besson CB, Lousada JL, Gaspar MJ, et al. (2016) Effects of Recent Minimum Temperature and Water Deficit Increases on *Pinus pinaster* Radial Growth and Wood Density in Southern Portugal. Frontiers in Plant Science 7.

Lachmuth S, Capblancq T, Keller SR, Fitzpatrick MC (2023) Assessing uncertainty in genomic offset forecasts from landscape genomic models (and implications for restoration and assisted migration). Frontiers in Ecology and Evolution 11.

Lazic D, Geßner C, Liepe KJ, et al. (2024) Genomic variation of European beech reveals signals of local adaptation despite high levels of phenotypic plasticity. Nature Communications 15:8553.

Leimu R, Fischer M (2008) A Meta-Analysis of Local Adaptation in Plants. PLOS ONE 3:e4010.

Long Z, Sang Y, Feng J, et al (2025) Evolutionary Genomics Unravels the Responses and Adaptation to Climate Change in a Key Alpine Forest Tree Species. Molecular Biology and Evolution 42:msaf116.

Marchi M, Bucci G, Iovieno P, Ray D (2024) ClimateDT: A Global Scale-Free Dynamic Downscaling Portal for Historic and Future Climate Data. Environments 11:82.

Mathur S, DeWoody JA (2021) Genetic load has potential in large populations but is realized in small inbred populations. Evolutionary Applications 14:1540–1557.

Matuszewski S, Hermisson J, Kopp M (2015) Catch Me if You Can: Adaptation from Standing Genetic Variation to a Moving Phenotypic Optimum. Genetics 200:1255–1274.

Mazza G, Cutini A, Manetti MC (2014) Influence of tree density on climate-growth relationships in a *Pinus pinaster* Ait. forest in the northern mountains of Sardinia (Italy). iForest - Biogeosciences and Forestry 8:456.

Messer PW, Petrov DA (2013) Population genomics of rapid adaptation by soft selective sweeps. Trends in Ecology and Evolution 28:10.1016/j.tree.2013.08.003.

Metheringham CL, Plumb WJ, Flynn WRM, et al. (2025) Rapid polygenic adaptation in a wild population of ash trees under a novel fungal epidemic. Science 388:1422–1425.

Metzger MJ (2018) The Environmental Stratification of Europe

Morando-Milà J, Grau O, Ulaszewski B, et al. (2026) Re-Interpreting Genetic Offset: Quantifying the Least Required Evolutionary Rate Under Climate Change at the Mediterranean Range Margin of European Beech. Evolutionary Applications 19:e70230.

Nadeau CP, Urban MC (2019) Eco-evolution on the edge during climate change. Ecography 42:1280–1297.

Niu S, Li J, Bo W, et al. (2022) The Chinese pine genome and methylome unveil key features of conifer evolution. Cell 185:204–217.e14.

Olsson S, Bouilli A, Niskanen A, et al. Range-wide SNP reference data and filtering effects in three major European pines. In preparation.

Olsson S, Grivet D, Westergren M, et al. (2025) Genetic Assignment at Different Geographical Levels: A Case Study in a Forest Tree Species (*Pinus pinaster* Ait.) Using SNP Markers. Evolutionary Applications 18:e70145.

Orr HA, Unckless RL (2014) The Population Genetics of Evolutionary Rescue. PLOS Genetics 10:e1004551.

Parmesan C (2006) Ecological and Evolutionary Responses to Recent Climate Change. Annual Review of Ecology, Evolution, and Systematics 37:637–669.

Rellstab C, Gugerli F, Eckert AJ, et al. (2015) A practical guide to environmental association analysis in landscape genomics. Molecular Ecology 24:4348–4370.

Saleh D, Chen J, Leplé J-C, et al. (2022) Genome-wide evolutionary response of European oaks during the Anthropocene. Evolution Letters 6:4–20.

Sánchez-Salguero R, Navarro-Cerrillo RM, Camarero JJ, Fernández-Cancio Á (2012) Selective drought-induced decline of pine species in southeastern Spain. Climatic Change 113:767–785.

Savolainen O, Pyhäjärvi T, Knürr T (2007) Gene Flow and Local Adaptation in Trees. Annual Review of Ecology, Evolution, and Systematics 38:595–619.

Scotti I, González-Martínez SC, Budde KB, Lalagüe H (2016) Fifty years of genetic studies: what to make of the large amounts of variation found within populations? Annals of Forest Science 73:69–75.

Shaw RG, Etterson JR (2012) Rapid climate change and the rate of adaptation: insight from experimental quantitative genetics. New Phytologist 195:752–765.

Shiogama H, Fujimori S, Hasegawa T, et al. (2023) Important distinctiveness of SSP3-7.0 for use in impact assessments. Nat Clim Chang 13:1276–1278.

Solé-Medina A, Hurel A, Avanzi C, et al. (2025) Macro- and micro-geographical genetic variation in early-fitness traits in populations of maritime pine (*Pinus pinaster*). Annals of Botany 135:655–668.

Taccoen A, Piedallu C, Seynave I, et al. (2019) Background mortality drivers of European tree species: climate change matters. Proceedings of the Royal Society B 286:20190386.

Theraroz A (2024) Maritime pine genetic resources: geographical variability, selection pressures and future adaptation. Theses, Université de Bordeaux.

Theraroz A, Guadaño-Peyrot C, Archambeau J, et al. (2024) The genetic consequences of population marginality: A case study in maritime pine. Diversity and Distributions 30:e13910.

Vekemans X, Hardy OJ (2004) New insights from fine-scale spatial genetic structure analyses in plant populations. Molecular Ecology 13:921–935.

Vendramin GG, Anzidei M, Madaghiele A, Bucci G (1998) Distribution of genetic diversity in *Pinus pinaster* Ait. as revealed by chloroplast microsatellites. Theoretical and Applied Genetics 97:456–463.

Wang J, Hill WG, Charlesworth D, Charlesworth B (1999) Dynamics of inbreeding depression due to deleterious mutations in small populations: mutation parameters and inbreeding rate. Genetics Research 74:165–178.

Yoder JB, Tiffin P (2018) Effects of Gene Action, Marker Density, and Timing of Selection on the Performance of Landscape Genomic Scans of Local Adaptation. Journal of Heredity 109:16–28.

Zelinka MD, Myers TA, McCoy DT, et al. (2020) Causes of Higher Climate Sensitivity in CMIP6 Models. Geophysical Research Letters 47:e2019GL085782.

