## Supplementary Material for "Rapid evolutionary responses to climate change in maritime pine (*Pinus pinaster* Ait.) despite widespread adaptive lag"

### *Supplementary Material (Methods)*

*SNP data filtering*

Quality control of the dataset 1 was performed following the criteria described in Francisco *et al.* (2026): CR ≥ 85, FLD ≥ 3.2, HetSO ≥ -0.3 and samples DQC ≥ 0.4 with QC call rate ≥ 85. In addition, monomorphic SNPs and variants with a minor allele count < 12 were removed, as well as SNPs and individuals with more than 30% missing data. This results in a final dataset comprising 8,469 SNPs from 1,518 trees across 82 populations.

*Climatic variable selection*

To identify climatic variables potentially driving adaptation, we followed Francisco *et al.* (2025) three-step procedure involving: (i) a pre-selection of climatic predictors based on the ecology of *P. pinaster*; (ii) a forward selection to identify the climatic predictors explaining the largest proportion of the genetic variance between populations; and (iii) the removal of highly correlated variables. The forward selection was achieved using the ordiR2step function from the VEGAN R package (Oksanen *et al.* 2025), from a null model with no climatic variables up to a full model incorporating all retained variables. Variables were retained if they increased the model's $R^2$, with significance assessed via 1,000 permutations and $p$-value < 0.05. Twenty runs were performed to ensure robustness of the forward selection. In the last step, climatic variables exhibiting high correlation (Pearson's $|r| > 0.75$) and/or variance inflation factors (VIF) over 10 (James *et al.* 2021) were removed, prioritising variables identified as the most informative in the forward selection step. Six climatic predictors were finally retained (See Main text).

*Mapping of SNPs onto a reference genome for genetic load annotation*

Annotation of SNPs to estimate realised and potential genetic load first involved mapping the SNPs onto the working version of the *Pinus tabuliformis* reference genome (Niu *et al.* 2022), one of the closest reference genomes currently available for *P. pinaster*. In this version, the 12 large chromosomes from the original reference genome (1.4 to 2.4 Gb in length; Niu *et al.* 2022) have been split, for a total of 22,739 contigs (Kesälahti *et al.* 2025). SNPs were mapped onto the *P. tabuliformis* genome by aligning their 120 bp flanking sequences using BLASTN (NCBI BLAST 2.2.26). The optimal alignment for each SNP was selected based on sequence identity, alignment length, mismatch count, and e-value. SNPs were also polarised in order to define the ancestral and alternative alleles, with the allele present in *P. tabuliformis* defined as ancestral.

*Climate-associated SNPs*

Climate-associated SNPs were identified using six gene-environment associations (GEA) methods:

Gradient forest (GF) is a tree-based machine learning framework designed to capture complex and, potentially non-linear relationships between environmental variables and allele frequency variation across populations (Ellis *et al.* 2012; Fitzpatrick *et al.* 2021). Analyses were conducted using either observed allele frequencies (GF-raw) or allele frequencies corrected for population genetic structure (GF-corrected). The corrected allele frequencies were obtained from LFMM-adjusted effect sizes combined with climatic predictors (Francisco *et al.* 2025). For each dataset, five independent GF models were fitted and the highest-ranking 1% of SNPs were retained. Climate-associated loci were defined as those consistently identified across all five replicates.

Associations between SNPs and climatic variables were also assessed using the standard covariate model implemented in BayPass (Gautier 2015). This Bayesian approach evaluates marker-environment relationships while accounting for shared demographic history through a population variance-covariance matrix. Loci exhibiting an average Bayes factor greater than 10 over five independent runs were retained, corresponding to “very strong evidence” for an association according to the Jeffreys' scale (Kass and Raftery 1995).

The latent factor mixed model (LFMM) approach was used as an additional genotype-environment association method. LFMM estimates the contribution of environmental predictors to genetic variation while simultaneously correcting for population genetic structure through latent factors (Caye *et al.* 2019). Two latent factors were included in the analyses, and loci with a false discovery rate (FDR)-adjusted significance level below 5% were considered significantly associated with the climatic predictors.

Finally, redundancy analysis (RDA) and partial redundancy analysis (pRDA) were applied to investigate multivariate relationships between genetic and environmental predictors. These constrained ordination methods identify combinations of environmental variables that explain genomic variation (Legendre and Legendre 2012). In pRDA, the influence of population genetic structure was accounted for by including the first two genomic principal components as covariates. Candidate loci were detected following the procedure described in Capblancq and Forester (2021), with SNPs showing extreme Mahalanobis distances along the first two canonical axes not exceeding a FDR of 5% identified as climate-associated SNPs.

### *Supplementary Material (Tables & Figures)*

*Table S1.* **Climatic models used at different time periods**. List of the five global climate models (GCMs) used to predict the historical (Mid-Holocene ~6,000 years ago) and future (2041-2070 period under the Socio-Economic Pathway 3-7.0) climates.

| Time period | Global climate models (GCMs) | Reference |
|---|---|---|
| Historical (Mid-Holocene ~6,000 years ago) | CNRM-CM5 | Voldoire *et al.* (2013) |
| | IPSL-CM5A-LR | Braconnot *et al.* (2013) |
| | MIROC-ESM | Watanabe *et al.* (2011) |
| | MPI-ESM-P | Giorgetta *et al.* (2013) |
| | MRI-CGCM3 | Yukimoto *et al.* (2012) |
| Future (2041-2070) | GFDL-ESM4.1 | Dunne *et al.* (2020) |
| | IPSL-CM6A-LR | Boucher *et al.* (2020) |
| | MPI-ESM1-2-HR | Gutjahr *et al.* (2019) |
| | MRI-ESM2-0 | Yukimoto *et al.* (2019) |
| | UKESM1-0-LL | Sellar *et al.* (2019) |

*Table S2.* **Simulation parameters**. Description of all the parameters used in the simulation framework.

| Parameter | Value |
|---|---|
| **Population size** | 10,000 |
| **Sex ratio** | 0.5 |
| **Number of chromosomes and chromosome size** | 12/1Mb |
| **Neutral mutation rates** | $1 \times 10^{-8}$ |
| **Recombination rate within chromosomes** | $1 \times 10^{-6}$ |
| **Number of offspring per female** | X~Poisson(λ=2) |
| **Seed dispersal (Lacanau/Tocchi)** | X~ Normal(0, σ=0.09/0.03) |
| **Pollen dispersal (Lacanau/Tocchi)** | Gaussian kernel with σ=0.30/0.15 |
| **Maximum pollen dispersal distance (Lacanau/Tocchi)** | 0.7/0.5 of the entire space |

*Table S3.* **Statistical tests**. Summary of the statistical tests and models used for each main analysis, alongside the corresponding test statistics, degrees of freedom (df), 95% confidence intervals (CI) and associated *p*-values.

| Variable | Test | Comparison | Statistic | df | 95% CI | *p*-value |
|---|---|---|---|---|---|---|
| **Genetic diversity ($H_i$)** | Two-sample Welch's *t*-test | $H_i$ ~ populations | $t$= 47 | 1467 | [0.053; 0.057] | < $2.2e^{-16}$ |
| | Welch's one-way Anova | $H_i$ ~ cohorts(populations) | $F$= 1002 | 593 | - | < $2.2e^{-16}$ |
| | Welch's one-way Anova Games-Howell post-hoc pairwise comparisons | Lacanau adult/ Lacanau juvenile | - | - | [0.018; 0.029] | $7.5e^{-13}$ |
| | | Lacanau adult/ Tocchi adult | - | - | [-0.012; -0.047] | $6.0e^{-14}$ |
| | | Lacanau adult/ Tocchi juvenile | - | - | [-0.046; -0.038] | $1.0e^{-12}$ |
| | | Lacanau juvenile/ Tocchi adult | - | - | [-0.079; -0.068] | 0 |
| | | Lacanau juvenile/ Tocchi juvenile | - | - | [-0.073; -0.060] | $3.3e^{-11}$ |
| | | Tocchi adult/ Tocchi juvenile | - | - | [0.003; 0.012] | $1.0e^{-5}$ |

| | | | | | | |
|---|---|---|---|---|---|---|
| **Realised genetic load** | Two-sample Welch's *t*-test | Realised load ~ populations | *t*= -40 | 1470 | [-17.95; -16.28] | < 2.2e$^{-16}$ |
| | Welch's one-way Anova | Realised load ~ cohorts(populations) | *F*= 612 | 651 | - | < 2.2e$^{-16}$ |
| | Welch's one-way Anova Games-Howell post-hoc pairwise comparisons | Lacanau adult/ Lacanau juvenile | - | - | [-9.60; -6.24] | 0 |
| | | Lacanau adult/ Tocchi adult | - | - | [13.66; 16.07] | 0 |
| | | Lacanau adult/ Tocchi juvenile | - | - | [12.17; 15.28] | 0 |
| | | Lacanau juvenile/ Tocchi adult | - | - | [21.07; 24.51] | 0 |
| | | Lacanau juvenile/ Tocchi juvenile | - | - | [19.68; 23.62] | 8.7e$^{-11}$ |
| | | Tocchi adult/ Tocchi juvenile | - | - | [-2.73; 0.45] | 0.254 |

| | | | | | | |
|---|---|---|---|---|---|---|
| **Potential genetic load** | Two-sample Welch's *t*-test | Potential load ~ populations | *t*= 57 | 1478 | [28.98; 31.04] | < 2.2e$^{-16}$ |
| | Welch's one-way Anova | Potential load ~ cohorts(populations) | *F*= 1169 | 637 | - | < 2.2e$^{-16}$ |
| | Welch's one-way Anova Games-Howell post hoc pairwise comparisons | Lacanau adult/ Lacanau juvenile | - | - | [4.24; 8.56] | 0 |
| | | Lacanau adult/ Tocchi adult | - | - | [-30.00; -27.04] | 4.4e$^{-13}$ |
| | | Lacanau adult/ Tocchi juvenile | - | - | [-28.67; -24.61] | 0 |
| | | Lacanau juvenile/ Tocchi adult | - | - | [-37.15; -32.69] | 0 |
| | | Lacanau juvenile/ Tocchi juvenile | - | - | [-35.66; -30.42] | 9.7e$^{-11}$ |
| | | Tocchi adult/ Tocchi juvenile | - | - | [-0.22; 3.98] | 0.10 |

| | | | | | | |
|---|---|---|---|---|---|---|
| **Climate (mal) adaptation RDA values** | Two-sample Welch's *t*-test | Lacanau adult/ Lacanau juvenile RDA1 | *t*= 0.168 | 542 | [-0.003; 0.003] | 0.867 |
| | | Lacanau adult/ Lacanau juvenile RDA2 | *t*= 2.045 | 592 | [0; 0.005] | 0.041 |
| | | Tocchi adult/ Tocchi juvenile RDA1 | *t*= -2.671 | 331 | [-0.010; -0.001] | 0.008 |
| | | Tocchi adult/ Tocchi juvenile RDA2 | *t*= -1.746 | 408 | [-0.006; 0] | 0.082 |

*Table S4.* **Wright's neighbourhood size (*Nb*)**. Estimates of *Nb* across populations for each of the ten replicates based on 500 randomly sampled SNPs and 1,000 permutations per replicate.

| **Run** | $Nb_{Lacanau}$ | $Nb_{Tocchi}$ |
|---|---|---|
| **1** | 774 | 109 |
| **2** | 693 | 115 |
| **3** | 820 | 90 |
| **4** | 546 | 97 |
| **5** | 664 | 102 |
| **6** | 560 | 106 |
| **7** | 648 | 122 |
| **8** | 727 | 104 |
| **9** | 629 | 102 |
| **10** | 559 | 112 |

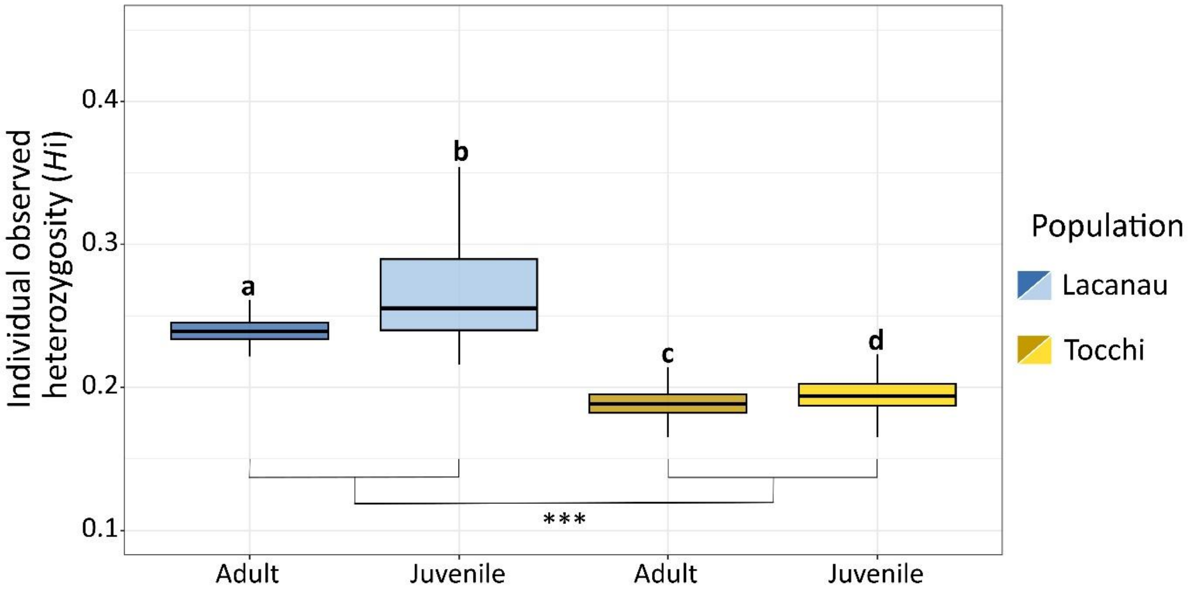


*Figure S1.* **Individual observed heterozygosity ($H_i$) across populations and cohorts**. Comparison of $H_i$ across populations and cohorts. Games-Howell's post hoc groups are shown by lowercase letters (*p*-value < 0.05). Pairwise population differences were tested using two-sample *t*-tests (***: *p*-value < 0.001).

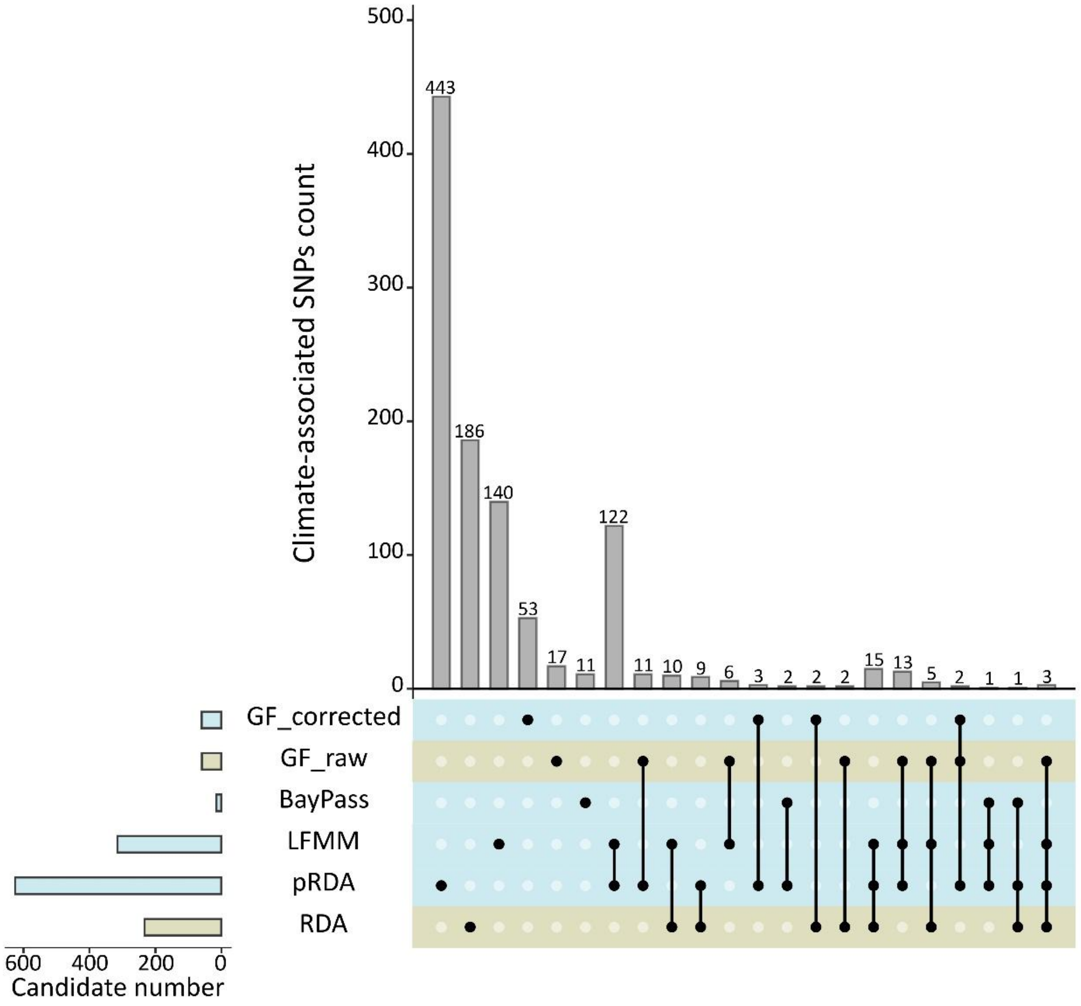


*Figure S2.* **Climate-associated SNPs identified by GEA methods**. UpSet plot summarising the climate-associated SNPs identified by each GEA method (horizontal bars), as well as information on whether the SNP identification is unique or shared across methods (vertical bars) with corresponding method(s) indicated by the dots below the vertical bars. Colours indicate whether the method corrects (in blue) or not (in green) for population genetic structure.

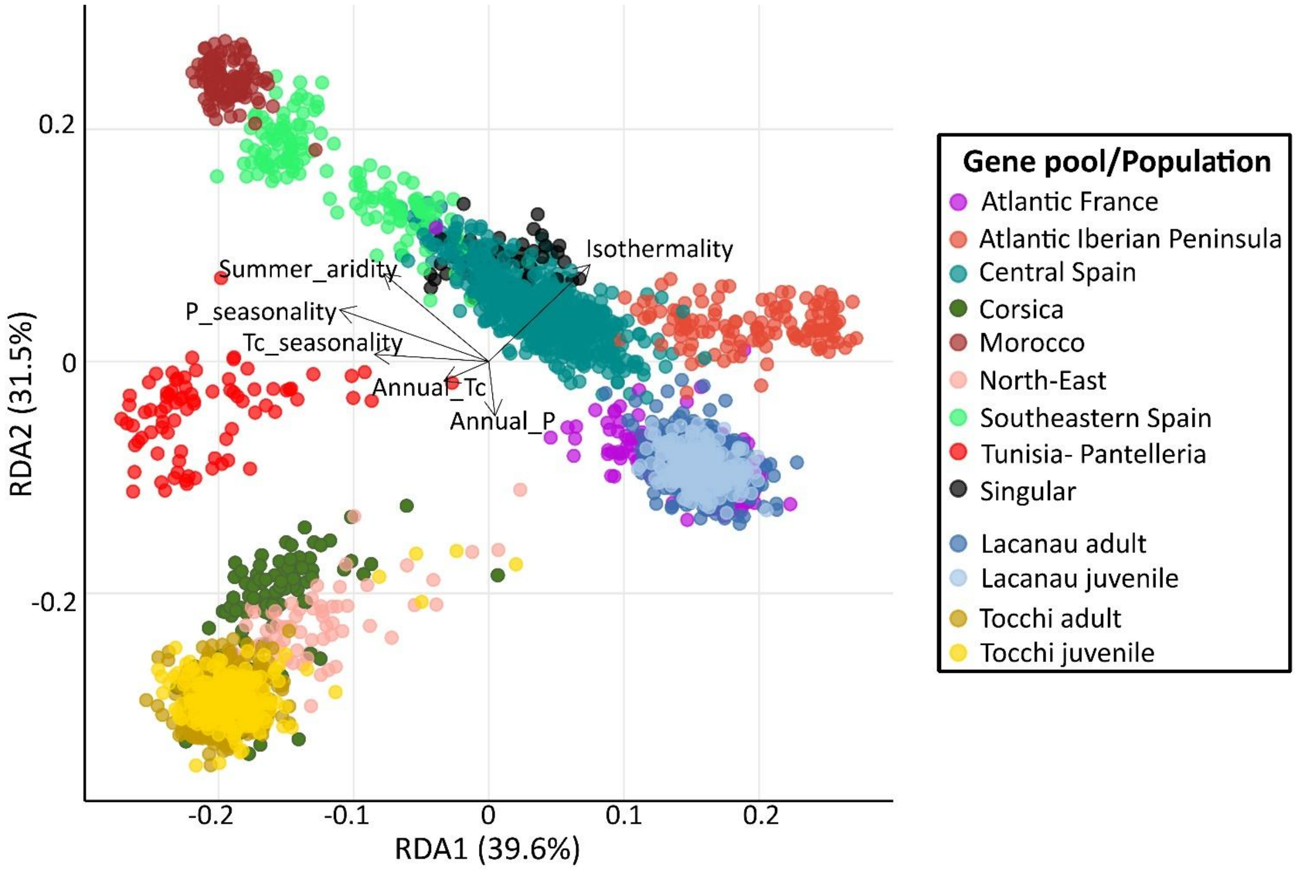


*Figure S3.* **Description of the RDA model used to infer climate (mal)adaptation**. Biplot of the RDA model showing the relative contribution of the climatic predictors to the first two RDA axes and the distribution of the individuals within populations used to fit the model. Genotyped individuals from Lacanau and Tocchi populations are projected into this space based on their genomic composition at climate-associated loci. Dots are coloured according to gene pool (Theraroz *et al.* 2024) or population cohort assignment. To facilitate readability, distinct gene pools consisting of a single population have been grouped under the term "*singular*" (see details in Theraroz *et al.* 2024).

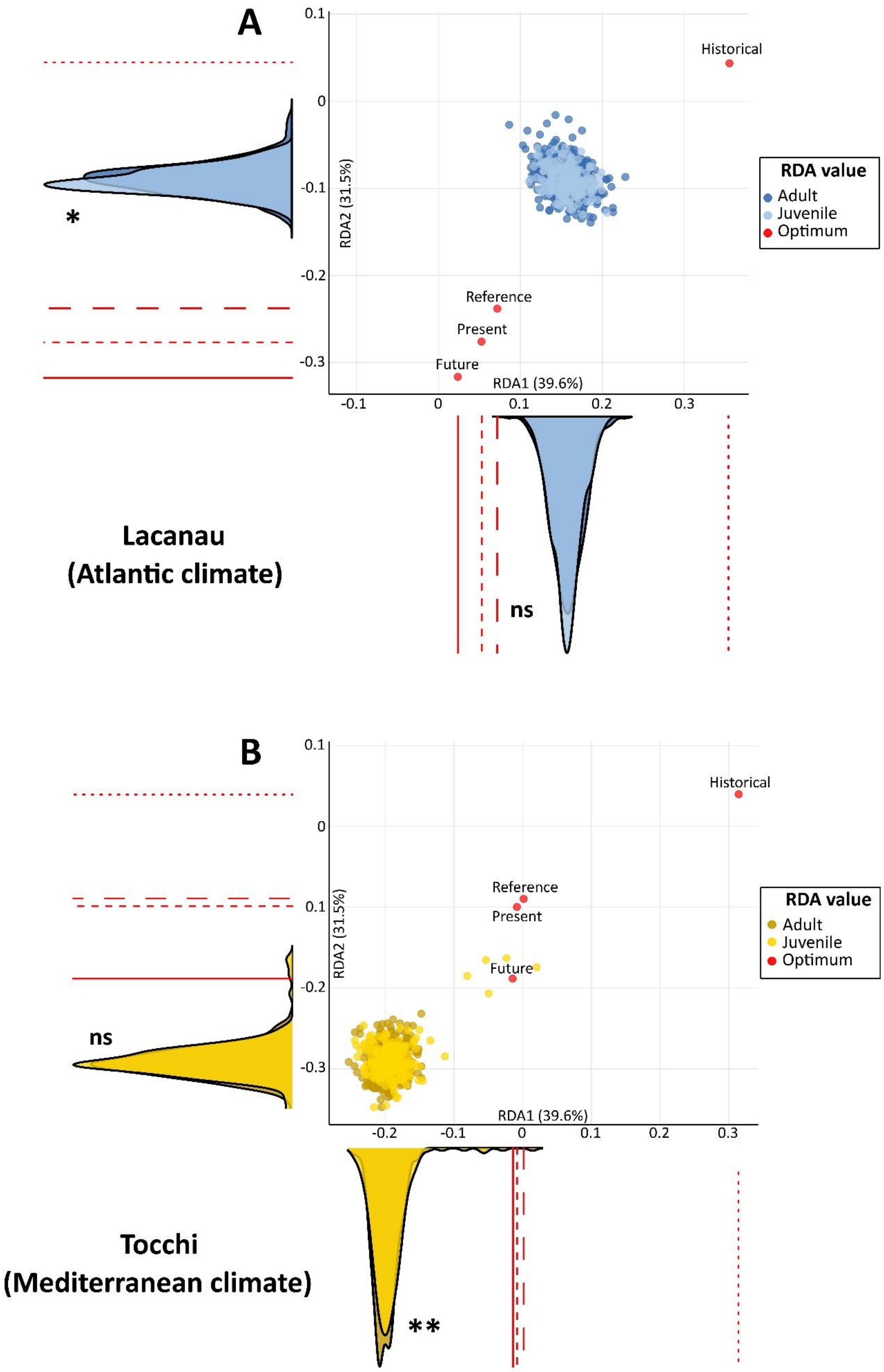
A
Historical
Reference
Present
Future
RDA1 (39.6%)
RDA2 (31.5%)
RDA value
Adult
Juvenile
Optimum
*
ns
Lacanau
(Atlantic climate)
B
Historical
Reference
Present
Future
RDA1 (39.6%)
RDA2 (31.5%)
RDA value
Adult
Juvenile
Optimum
ns
**
Tocchi
(Mediterranean climate)

*Figure S4.* **Within-population variability in climate (mal)adaptation and potential for rapid evolutionary responses including reference optimum**. Focus on the projected positions of genotyped individuals from (**A**) Lacanau and (**B**) Tocchi within the RDA ordination presented in Figure 2. Individuals are positioned according to their genomic composition at climate-associated loci and coloured by cohort, expressed as RDA scores. Predicted genomic composition optima, also expressed as RDA scores, are shown for historical (Mid-Holocene, approximately 6,000 years ago), reference (1901-1950), present (1991-2020) and future (2041-2070) time periods. Density plots show cohort-specific distributions of RDA scores, with differences between cohorts assessed using Welch's two-sample *t*-tests ($^{ns}$ not significant; *: *p*-value < 0.05; **: *p*-value < 0.01). Optimum values are also represented as lines on density plots, corresponding to historical (dotted), reference (long-dashed), present (short-dashed), and future (solid) time periods.

### *References*


Boucher O, Servonnat J, Albright AL, *et al.* (2020) Presentation and Evaluation of the IPSL-CM6A-LR Climate Model. *Journal of Advances in Modeling Earth Systems* 12:e2019MS002010.

Braconnot P, Denvil S, Foujols M-A, *et al.* (2013) IPSL-CM5A-LR model output prepared for CMIP5 midHolocene experiment.

Capblancq T, Forester BR (2021) Redundancy analysis: A Swiss Army Knife for landscape genomics. *Methods in Ecology and Evolution* 12:2298-2309.

Caye K, Jumentier B, Lepeule J, François O (2019) LFMM 2: Fast and Accurate Inference of Gene-Environment Associations in Genome-Wide Studies. *Molecular Biology and Evolution* 36:852-860.

Dunne JP, Horowitz LW, Adcroft AJ, *et al.* (2020) The GFDL Earth System Model Version 4.1 (GFDL-ESM 4.1): Overall Coupled Model Description and Simulation Characteristics. *Journal of Advances in Modeling Earth Systems* 12:e2019MS002015.

Ellis N, Smith SJ, Pitcher CR (2012) Gradient forests: calculating importance gradients on physical predictors. *Ecology* 93:156-168.

Fitzpatrick MC, Chhatre VE, Soolanayakanahally RY, Keller SR (2021) Experimental support for genomic prediction of climate maladaptation using the machine learning approach Gradient Forests. *Molecular Ecology Resources* 21:2749-2765.

Francisco T, Lesur-Kupin I, Guadaño-Peyrot C, *et al.* (2026) Demographic history shapes forest tree vulnerability to climate change. 2026.03.10.710859

Francisco T, Mayol M, Vajana E, *et al.* (2025) Genomic signatures of climate-driven (mal)adaptation in an iconic conifer, the English Yew (*Taxus baccata* L.). *Evolutionary Applications* 18:e70160.

Gautier M (2015) Genome-Wide scan for adaptive divergence and association with population-specific covariates. *Genetics* 201:1555-1579.

Giorgetta MA, Jungclaus J, Reick CH, *et al.* (2013) Climate and carbon cycle changes from 1850 to 2100 in MPI-ESM simulations for the Coupled Model Intercomparison Project phase 5. *Journal of Advances in Modeling Earth Systems* 5:572-597.

Gutjahr O, Putrasahan D, Lohmann K, *et al.* (2019) Max Planck Institute Earth System Model (MPI-ESM1.2) for the high-resolution model intercomparison project (HighResMIP). *Geoscientific Model Development* 12:3241-3281.

James G, Witten D, Hastie T, Tibshirani R (2021) An introduction to statistical learning: with applications in R. Springer US, New York, NY

Kass RE, Raftery AE (1995) Bayes Factors. *Journal of the American Statistical Association* 90:773-795.

Kesälahti R, Kumpula TA, Cervantes S, *et al.* (2025) Optimising exome captures in species with large genomes using species-specific repetitive DNA blocker. *Molecular Ecology Resources* 25:e14053.

Legendre P, Legendre L (2012) Chapter 11 - Canonical analysis. In: Legendre P, Legendre L (eds) *Developments in Environmental Modelling*. Elsevier, pp 625-710

Niu S, Li J, Bo W, *et al.* (2022) The Chinese pine genome and methylome unveil key features of conifer evolution. *Cell* 185:204-217.e14.

Oksanen J, Simpson GL, Blanchet FG, *et al.* (2025) vegan: Community Ecology Package. 2.6-8

Sellar AA, Jones CG, Mulcahy JP, *et al.* (2019) UKESM1: description and evaluation of the U.K. Earth system model. *Journal of Advances in Modeling Earth Systems* 11:4513-4558.

Theraroz A, Guadaño-Peyrot C, Archambeau J, *et al.* (2024) The genetic consequences of population marginality: A case study in maritime pine. *Diversity and Distributions* 30:e13910.

Voldoire A, Sanchez-Gomez E, Salas y Mélia D, *et al.* (2013) The CNRM-CM5.1 global climate model: description and basic evaluation. *Climate Dynamics* 40:2091-2121.

Watanabe S, Hajima T, Sudo K, *et al.* (2011) MIROC-ESM 2010: model description and basic results of CMIP5-20c3m experiments. *Geoscientific Model Development* 4:845-872.

Yukimoto S, Adachi Y, Hosaka M, *et al.* (2012) A new global climate model of the meteorological research institute: MRI-CGCM3 Model description and basic performance. *Journal of the Meteorological Society of Japan* Ser II 90A:23-64.

Yukimoto S, Kawai H, Koshiro T, *et al.* (2019) The meteorological research institute earth system model version 2.0, MRI-ESM2.0: Description and basic evaluation of the physical component. *Journal of the Meteorological Society of Japan* Ser II 97:931-965.
